# Inhibition of Lipin lipid phosphatase hyperactivity rescues TorsinA neurological disease

**DOI:** 10.1101/606947

**Authors:** Ana Cascalho, Joyce Foroozandeh, Lise Hennebel, Christine Klein, Stef Rous, Beatriz Dominguez Gonzalez, Antonio Pisani, Maria Meringolo, Sandra F. Gallego, Patrik Verstreken, Philip Seibler, Rose E. Goodchild

**Affiliations:** VIB-KU Leuven Center for Brain and Disease Research; Dept of Neurosciences, KU Leuven, 3000 Leuven, Belgium; Institute of Neurogenetics, University of Lübeck, Lübeck, Germany; Laboratory of Neurophysiology and Plasticity, IRCCS Fondazione Santa Lucia and Department of Systems Medicine, University Tor Vergata, Rome, Italy; Leuven Brain Institute, 3000 Leuven, Belgium

## Abstract

TOR1A/TorsinA mutations cause poorly explained neurological diseases. A dominantly inherited mutation causes isolated dystonia, while biallelic mutations cause a recessive infant-onset syndrome with cases of lethality. Here we report an unexpected connection between lipid metabolism and these diseases. Lipin phosphatidic acid phosphatase activity was abnormally regulated in TorsinA dystonia patient cells, and in the brains of three different TorsinA disease model mice. Lipin activity was causative to symptoms given that lowering *Lipin1 in vivo* strongly intervened against lethality in disease mice. Furthermore, Lipin hyperactivity caused cell death *in vitro*, and *Lipin1* deficiency suppressed neurodegeneration *in vivo*. In addition, it protected the striatal cholinergic interneurons that are implicated in TorsinA movement disorders, and concomitantly suppressed abnormal motor behaviors of TorsinA mice. These data establish the central role of Lipin lipid enzyme hyperactivity in TorsinA disease and show that Lipin inhibition is a therapeutic target for these incurable conditions.

**One Sentence Summary:** Lipin inhibition rescues TorsinA neurological disease

---

TOR1A/TorsinA mutations cause at least two neurological diseases. Dominantly inherited TOR1A/TorsinA dystonia (OMIM #128100) caused by the +/ΔE mutation is characterized by childhood-onset involuntary twisting movements and abnormal postures of unexplained origin (1). Indeed, despite many years of study, no structural or degenerative pathology has been found to explain this isolated dystonia. It is the most common hereditary dystonia, and the genetic insult is a widely used experimental tool to investigate mechanisms leading to dystonia. However, it also remains poorly understood at molecular, cellular and neurobiology system levels. There is also a recessive TorsinA syndrome with a broader set of symptoms. This emerges at birth, and is caused by TorsinA ΔE/ΔE or other biallelic combinations of TorsinA deletions (2–5). Affected infants invariably show arthrogryposis (joint contractures at birth), tremor, and develop speech, cognition and motor deficits.

There is no cure for either TorsinA disease. Dominant TorsinA dystonia symptoms often develop into severe, generalized dystonia that is highly debilitating and interferes with all aspects of daily life. It is typically managed with deep brain stimulation. However, this is invasive and often fails to provide full symptom relief. The alternative is botulinum toxin treatment of affected muscle groups if the dystonia remains focal or segmental, but this is also purely symptomatic and only partially effective. There are even fewer options for recessive syndrome patients, and there are examples where infants failed to survive even in high-level health care settings. The surviving children continue to suffer from severe neurological defects, including intellectual impairment, and require continuous support and care (2–5).

Poor understanding of disease molecular and cellular pathology is rate-limiting for translating information on the genetic cause into therapies. TorsinA is an ATPase that resides in the endoplasmic reticulum (ER) (6, 7). In turn, the ER operates in membrane protein quality control, calcium buffering, and lipid synthesis, while specialized ER domains like the nuclear envelope have additional functions. Torsin AAA+ enzymes have been shown to affect the three core ER functions, as well as several at the nuclear envelope, with the nature and severity of torsin loss-of-function phenotypes varying between cell types and experimental paradigms (8–14). This is in fact explainable by the inter-dependent nature of different ER functions (15). However, unfortunately, it means we lack information on which defects are primary to TorsinA mutation(s) and, more importantly, which are causative for disease and thus should be targeted for therapy.

The ability to dissect disease cause(s) has been particularly challenging because TorsinA diseases are defined by their behavioral symptoms. Thus, any suspected cause must be interrogated against behavioral abnormalities. This is difficult since animal models appear to tolerate the dominant ΔE/+ mutation that causes dystonia in humans, while mice with recessive syndrome mutations (−/−, −/ΔE and ΔE/ΔE knock-in) die as neonates (Figure 1A) (16, 17). More recently, a conditional TorsinA mouse model was developed that has features of both diseases. This carries one copy of TorsinA ΔE and one floxed TorsinA allele. *In vivo* expression of Cre-recombinase reduces the flox/ΔE genotype to biallelic −/ΔE: a recessive syndrome genotype. However, for as yet undetermined reasons, Cre expression during nervous system appears to most strongly impact animal movement, including causing dystonia-like symptoms reminiscent of those in TorsinA dystonia patients (18, 19). Here we refer to this model as the “cKO/ΔE” mouse.

**Fig. 1.**
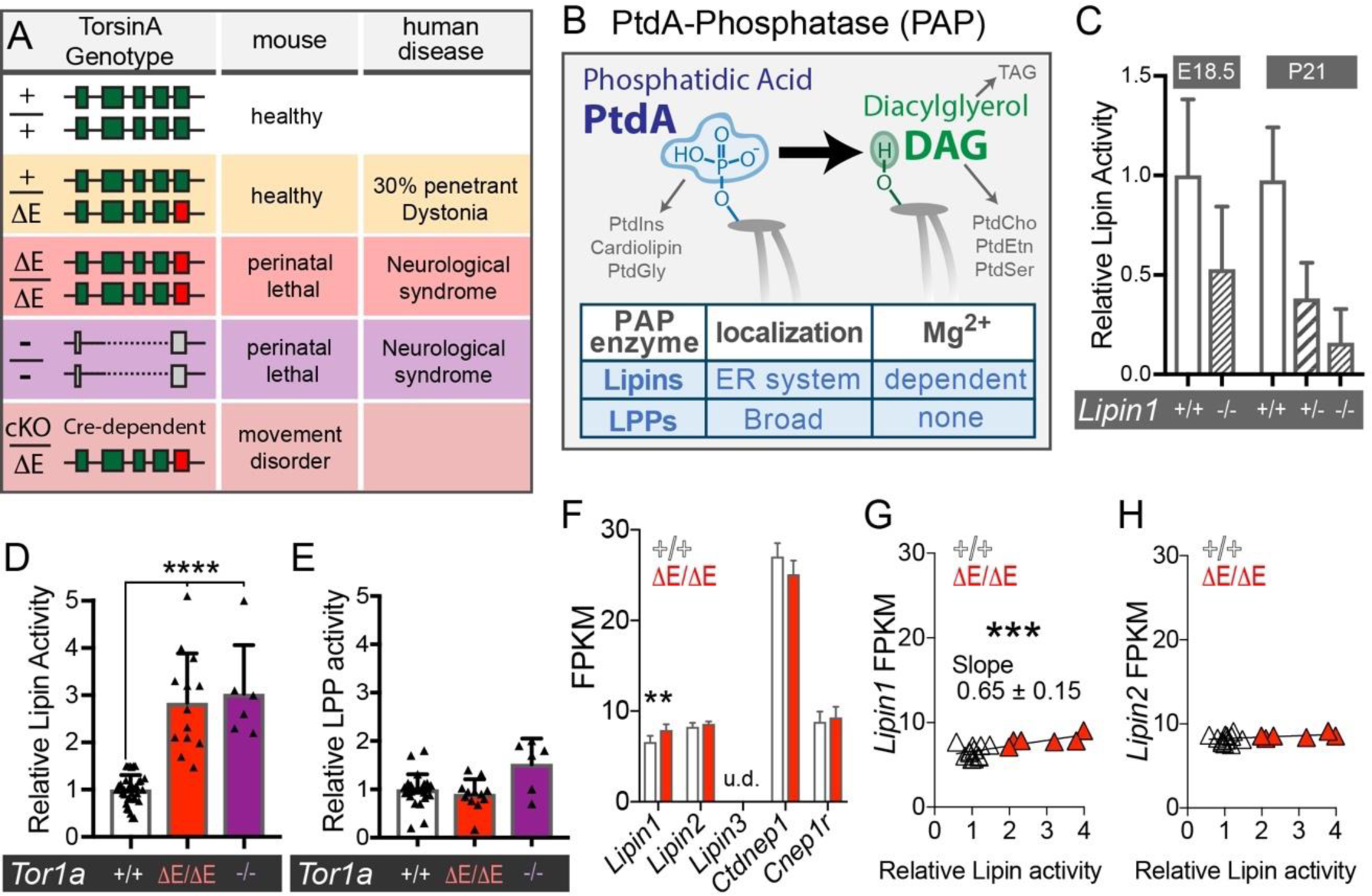
Lipin PAP activity is elevated in TorsinA disease mice. A) TorsinA genotypes, mouse phenotypes and human disease stages. B) Cartoon depicting PtdA conversion to DAG and derived lipid products. C – E) PAP activity in embryonic day 18.5 (E18.5) brain lysates relative to the TorsinA +/+ mean. (C) Confirmation that the assay detects Lipin PAP activity using *Lipin1*^−/+^ and *Lipin1*^−/−^. (D) Increased Lipin PAP activity in ΔE/ΔE *and* −/− brain; **** indicates significant difference (p<0.0001; One-way ANOVA). Points show values from individual embryos and bars show mean and standard deviation (SD). (E) LPP PAP activity is unaffected by *Tor1a* genotype. LPP activity is defined as the magnesium-independent fraction of PAP activity present in E18.5 brain lysates. (F) Lipin and Lipin-regulator gene expression as Fragments per Kilobase Million (FPKM) from RNAseq analysis of E18.5 TorsinA +/+ (n = 12) and ΔE/ΔE (n = 6) brain. u.d. refers to undetectable *Lipin3* mRNA. Bars show mean +/− SD. ** indicates significant difference (p < 0.01; One-way ANOVA). (G & H) Linear regression correlates brain *Lipin1* and *Lipin2* expression against relative Lipin PAP activity in individual +/+ and ΔE/ΔE embryos. The slope of *Lipin1* FPKM/ PAP activity significantly differs from zero (p = 0.004)

Here we use three TorsinA mouse models (−/−, ΔE/ΔE and cKO/ΔE) to examine the mammalian and nervous system relevance of our previous finding that *Drosophila dTorsin* regulates lipid synthesis. We focused on the hypothesized underlying mechanism in flies: regulation of phosphatidic acid (PtdA) dephosphorylation (12). This enzymatic reaction is mediated by Lipin or LPP enzymes, is a key step in *de novo* lipid synthesis, and produces the DAG signaling lipid (Figure 1B) (15). We now show that the Lipin phosphatidic acid phosphatase (PAP) enzyme is hyperactive in TorsinA −/−, ΔE/ΔE, and cKO/ΔE mouse brains. TorsinA also synergizes with TorsinB to regulate Lipin, and this underlies why a non-neuronal TorsinA cell line has normal Lipin activity. In line with this, knock down of TorsinB reveals abnormal Lipin activity in non-neuronal human patient cells with TorsinA mutations. Moreover, we establish *in vivo* that Lipin has a pathogenic role in TorsinA-associated neurodegeneration, motor dysfunction and animal lethality, and see that these phenotypes are rescuable by even partial reduction in Lipin hyperactivity. In conclusion, these data establish that Lipin hyperactivity is key in TorsinA neurological diseases, and that Lipin inhibition has strong therapeutic potential for these currently incurable conditions.

## Results

We previously demonstrated that fly *dTorsin* loss affects lipid synthesis in the larval fat body (12). However, another study failed to find lipidomic defects in mammalian cells (20). We now compared the brain lipidome of embryonic day 18.5 (E18.5) wild-type versus TorsinA ΔE/ΔE, or −/− littermates. This also revealed similar bulk steady-state glycerophospholipid levels (Supplemental Figure 1A – F). However, steady-state lipidomics lacks resolution to detect transient and/ or localized differences in lipid levels, particularly for low abundance lipids. It also provides little information on the rate of lipid enzyme reactions. We therefore specifically examined the metabolic reaction most effected by fly *dTorsin* loss: PtdA dephosphorylation to DAG (Figure 1B). Animals have two classes of PAP enzymes: Lipins in the ER, and Lipid-phosphate phosphatases (LPPs) in other regions of the cell (Figure 1B) (15). We confirmed both activities were present in embryonic brain lysates (Figure 1C, Supplemental Figure 1G) and examined whether either was affected by recessive TorsinA disease mutations. This revealed ∼3-fold elevated Lipin-mediated PAP activity in ΔE/ΔE and −/− brains compared to wild-types of the same litters (Figure 1D). In contrast, LPP enzyme activity was normal (Figure 1E). Thus, TorsinA indeed regulates brain lipid metabolism, and this function is affected by disease-causing genotypes.

Lipin also operates as a cofactor of PPAR transcription factors (21). We examined whether TorsinA also regulates this function by profiling gene expression in TorsinA mutant mouse brains. This found that PPAR genes and their targets were similarly expressed in +/+ and ΔE/ΔE brain (Supplemental Figure 2 and 3). We also examined Lipin gene expression to determine whether elevated enzyme activity originated from elevated transcription. Of the 3 mammalian Lipin genes, *Lipin1* and *Lipin2* transcripts were detected in embryonic brains (Figure 1F). *Lipin1* expression was mildly elevated in the ΔE/ΔE brain compared to wild-types (Figure 1F), and positively correlated with elevated enzyme activity (Figure 1G & H). However, *Lipin1* expression levels overlapped substantially between wild-type and ΔE/ΔE embryos, and differences were minor compared to altered enzyme activity. We also examined expression of the Lipin regulatory proteins, *Ctdnep1* and *Cnep1r1* (22, 23), and again saw similar expression in wild-type and ΔE/ΔE TorsinA mutant brain (Figure 1F). Thus, it appears that TorsinA regulates brain Lipin PAP activity without altering its co-transcriptional function.

We next asked whether excess Lipin activity contributes to disease. We approached this *in vivo* in cKO/ΔE mice where Nestin-Cre creates the KO allele during nervous system development (Figure 1A and 2A). This is shown to produce animals that survive past birth and develop behavioral impairments like those of dominant and recessive TorsinA disease patients (18). Similar to lipin activity in ΔE/ΔE and −/− brains, we again confirmed Lipin was hyperactive in cKO/ΔE mice: at E18.5 their brain Lipin activity was ∼2-fold higher than littermate controls (Figure 2B). We then modulated cKO/ΔE Lipin PAP activity by genetically reducing *Lipin1* expression (Figure 2B) (24). Complete *Lipin1/LIPIN1* loss is lethal in mouse and causes muscle disease in human (24–26), and thus we focused on physiologically tolerated *Lipin1*+/−. This was sufficient to reduce cKO/ΔE brain Lipin hyperactivity, although it was still significantly higher than that of controls.

**Figure 2.**
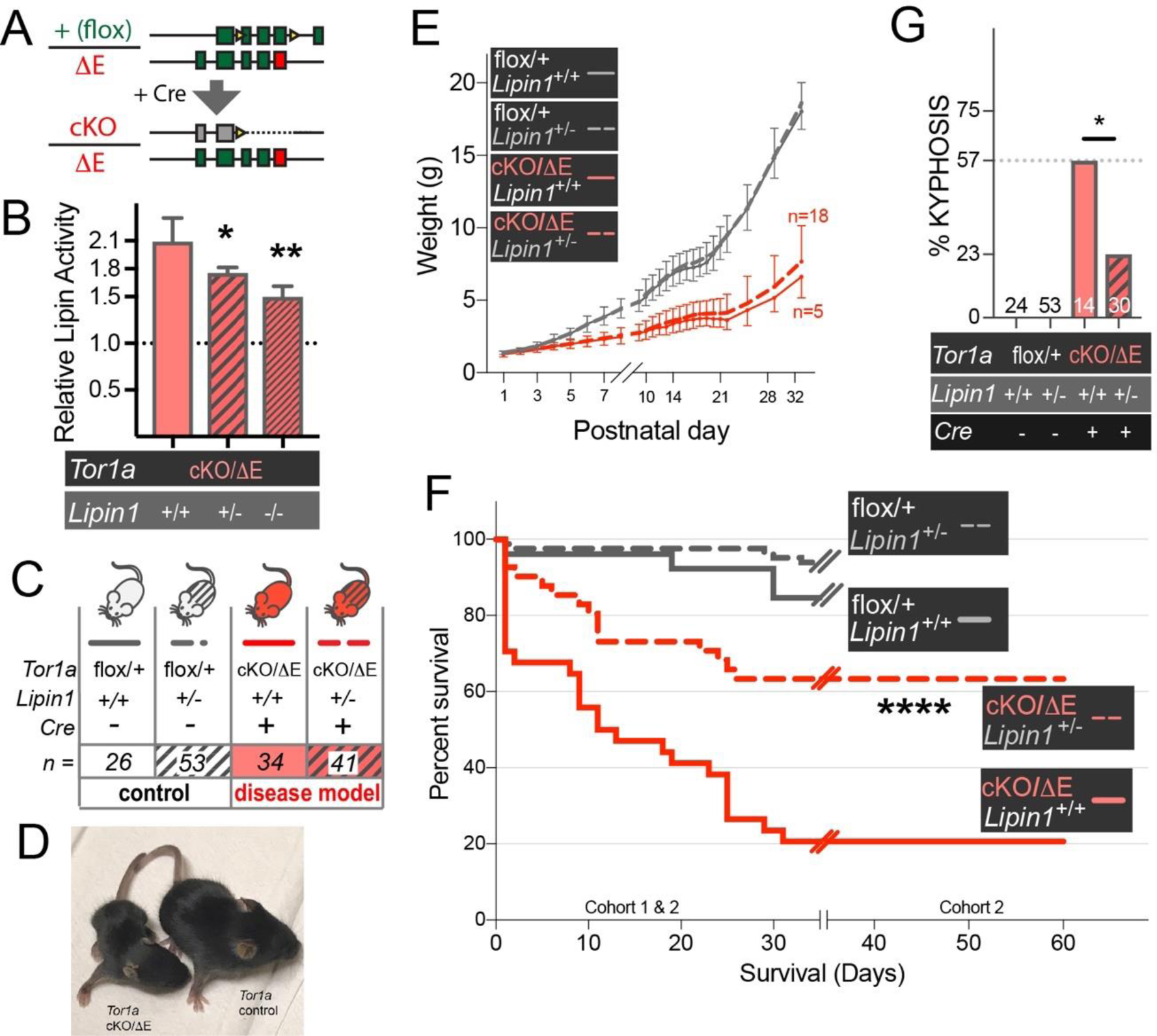
Lipin inhibition rescues TorsinA disease mouse lethality. A) Nestin-Cre deletion of TorsinA combined with the “ΔE” allele creates the cKO/ΔE TorsinA disease model mouse. B) Brain Lipin PAP activity is elevated in E18.5 cKO/ΔE, and partially rescued by *Lipin1* deletion. n = 4. *, ** (p = 0.016, 0.002; One-way ANOVA). C) Experimental series to test whether Lipin hyperactivity contributes to cKO/ΔE neurological dysfunction. D & E) cKO/ΔE animals fail to thrive. (D) cKO/ΔE mouse is significantly smaller than TorsinA wild-type (flox/+). (E) cKO/ΔE mice have normal birth weight but fall behind control littermates (grey line) independent of *Lipin* genotype (red lines). Numbers in the legend show the number of mice at birth, while numbers on right show cKO/ΔE number surviving until 30 days. F) Lipin1 deficiency strongly rescues lethality of cKO/ΔE mice. Lines show survival of TorsinA flox/+ and cKO/ΔE that are *Lipin1*+/+ or *Lipin1*+/−. Data from 0-30 days comes from two cohorts (*n* shown in C), while 30-60 days comes from a single cohort. **** significant (p < 0.0001) difference between cKO/ΔE:*Lipin1*+/+ and cKO/ΔE:*Lipin1*+/− (Log-rank Mantel-Cox test). G) Kyphosis frequency is significantly higher in cKO/ΔE compared with TorsinA flox/+ and * indicates a significant reduction in cKO/ΔE:*Lipin1*+/− (p = 0.027; Two-tailed Chi-square). Values show the number of mice surviving to P21 testing point.

We proceeded to test whether Lipin hyperactivity contributed to neurological defects in the cKO/ΔE mouse model (Figure 2C). Our first observation was that cKO/ΔE animals were more severely affected than previously reported. While their weight was normal at birth, they immediately failed to thrive (Figure 2D & E) and only ∼20% survived to postnatal day 30 (P30; Figure 2F). This is in fact consistent with other studies where Nestin-Cre drives a TorsinA loss-of-function mutation (27). It also mimics the symptoms of recessive TorsinA neonates who require immediate medical intervention and a proportion fail to survive (2–5). Strikingly, *Lipin1*+/− very strongly promoted cKO/ΔE survival (Figure 2F). Indeed, the majority of cKO/ΔE survived if they were deficient for *Lipin1* (Figure 2F). Further, cKO/ΔE animals stabilized once they survived to P30, and thus *Lipin1*+/− deficiency appeared to provide a permanent benefit (Figure 2F). Examination of kyphosis (hunchback) as a proxy for health of juvenile cKO/ΔE mice also detected a benefit of *Lipin1*+/− loss (Figure 2G). This was significant even though almost double the number of “uncorrected” cKO/ΔE mice had succumbed by this time point (53% and 27% loss of cKO/ΔE with *Lipin1*+/+ vs. *Lipin1*+/−, respectively). Thus, Lipin hyperactivity arising from TorsinA loss indeed plays a major role in the behavioral defects related to TorsinA recessive disease, and mild Lipin deficiency is sufficient to strongly improve survival of disease model mice.

The neurological defects of cKO/ΔE mice occur alongside neurodegeneration (18, 19) and we examined whether Lipin is sufficient to cause cell death in cultured cells. We examined wild-type mouse Lipin1 overexpression, as well as Lipin1 that is catalytically dead for PAP activity. In addition, Lipin1 undergoes multi-site phosphorylation that inhibits activity and targets Lipin1 protein for degradation. We therefore included mutants where 17 Ser/Thr residues were exchanged to Ala and thus blocked inhibitory phosphorylation events (29, 30). The four Lipin1 cDNAs were electroporated, and cell survival assayed the following day. This identified that Lipin1 overexpression is cytotoxic when phosphorylatable residues are mutated, and a large proportion of this relates to Lipin catalytic activity (Figure 3A). We then examined whether Lipin hyperactivity contributes to cell death *in vivo* in cKO/ΔE animals. We chose a juvenile developmental stage (P15) given the lethality of cKO/ΔE animals with uncorrected Lipin (Figure 2F). We used activated caspase labeling to detect apoptotic cells and, since we did not detect obvious hotspots of apoptosis, quantified staining across the entire brain. This indeed detected elevated apoptotic staining in P15 cKO/ΔE brain compared to wild-type or *Lipin1*+/− mice (Figure 3B). The elevated apoptotic labeling was significantly lower in “rescued” cKO/ΔE:*Lipin1*+/− mice.

**Fig. 3.**
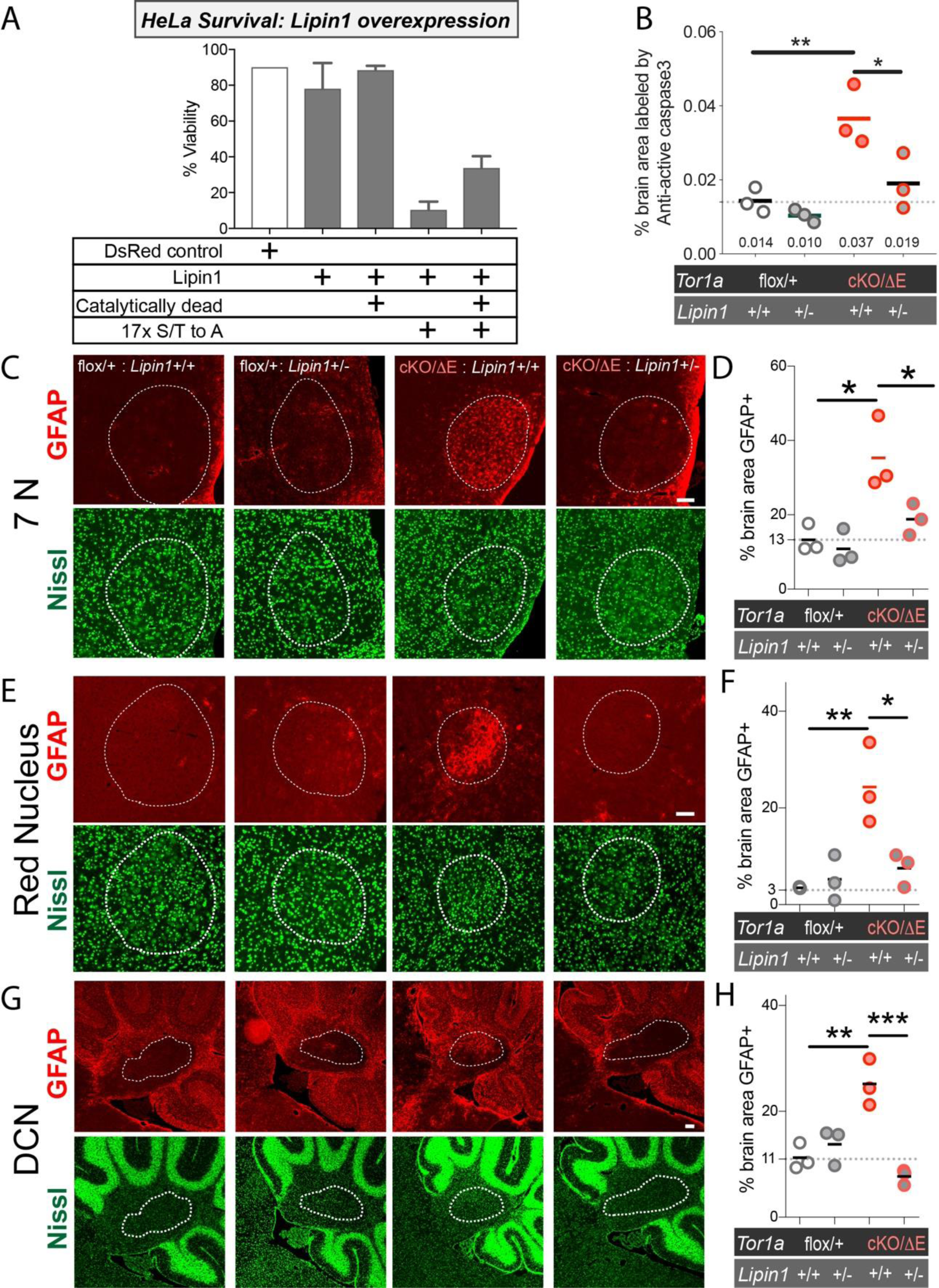
Lipin hyperactivity causes cell death *in vitro* and *in vivo* in cKO/ΔE. A) Viability of HeLa cells one-day after electroporation with dsRed-ER control (white bar), or mouse Lipin1 cDNAs. Bars show mean +/− SEM from 3 independent experiments. B) Area of activated caspase3 immunolabeling across entire hemispheres of P15 mice. Values present the mean for each group. *, ** indicate that cKO/ΔE:Lipin1+/+ significantly differ from controls or cKO/ΔE:Lipin1+/− (p < 0.05, 0.01. Two-Way ANOVA). C – H) GFAP staining detects reactive gliosis in cKO/ΔE that is rescued by *Lipin1* deficiency. (C, E, G) Images from P14 brain of anti-GFAP (red) and florescent Nissl staining (green). White dashed lines delineate nuclei of interest as identified by Nissl staining. Scale bars show 20 μm. (D, F, H) Quantitation of reactive gliosis as the % of each brain nucleus occupied by GFAP labeling. Points show value from each animal, and bars show the mean of the group *, **, *** indicate significant differences between genotypes (p < 0.05, 0.01, 0.001. 2-Way ANOVA).

We then examined reactive gliosis as a second measure of *in vivo* degeneration. We focused on three brain nuclei where we confirmed previous reports that anti-GFAP detects this event in cKO/ΔE (Nestin-Cre) animals; the facial nucleus (7N) (Figure 3C & 3D), red nucleus (Figure 3E & 3F), and deep cerebellar nuclei (Figure 3G & 3H) (18). This labeling and quantification showed that Lipin deficiency strongly rescued reactive gliosis. In fact, GFAP labeling was no different to wild-type animals. From these data we conclude that Lipin hyperactivity is a major contributor to the lethality and associated neurodegeneration arising from biallelic TorsinA mutations.

cKO/ΔE animals are also regarded as a phenotypic model for dystonia since neurodegeneration is particularly severe in motor circuits, including striatal cholinergic interneurons that are strongly implicated in dystonia pathogenesis (18, 19, 31). We now confirmed reports showing that regions of cKO/ΔE striatum have under half the normal number of cholinergic cells. (Figure 4A vs. 4C; Figure 4E red lines). As expected, cholinergic neuronal number was normal in *Lipin1+/−* mice (Figure 4A vs. 4B; 4E grey lines). We then examined “rescued” cKO/ΔE:*Lipin1+/−* mice. Qualitative analysis of anti – choline acetyltransferase (ChAT) labeling showed relatively normal signal (Figure 4A vs 4D). This was confirmed by quantitation detecting a significantly higher number of cholinergic neurons in cKO/ΔE:*Lipin1+/−* striatum than cKO/ΔE:*Lipin1+/+* (Figure 4E; compare red lines).

**Figure 4.**
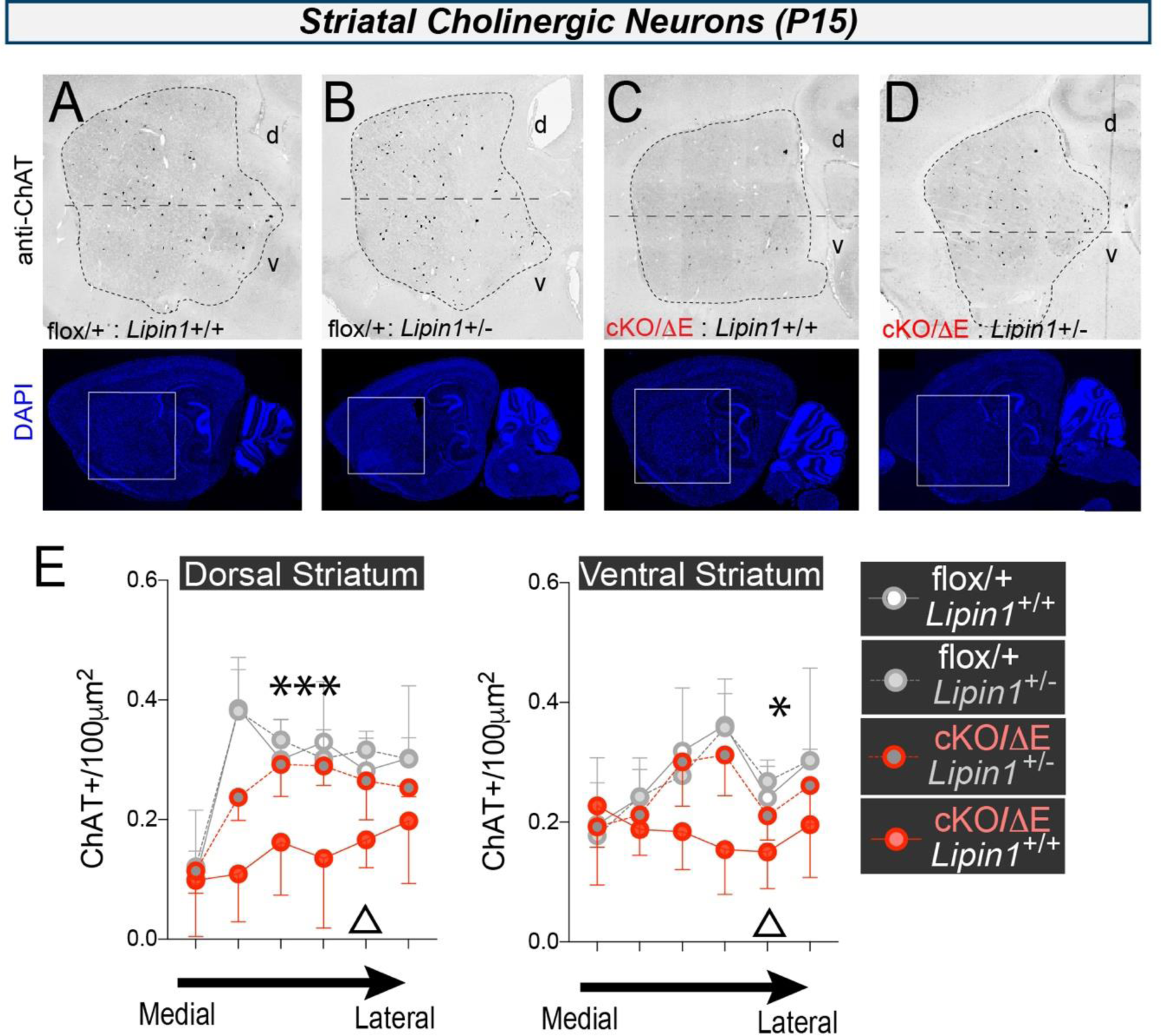
Lipin hyperactivity contributes to the sensitivity of cKO/ΔE striatal cholinergic interneurons. A – D) Representative images of Choline Acetyltransferase (ChAT) positive cells in the striatum of sagittal sections through the P15 mouse brain. DAPI shows the overview. d (dorsal), v (ventral). n = 3 animals/group. Anti-ChAT images were inverted to more clearly show staining of individual neurons; all images were identically treated. E) Cholinergic neuron density in dorsal and ventral striatum in sets of sections binned by mediolateral position; triangles show position of sections in A – D. ***, * reflect significant difference in cholinergic cell density between cKO/ΔE:*Lipin1*+/+ and cKO/ΔE:*Lipin1*+/−. p = 0.0028; p = 0.015; Two-way ANOVA of dorsal and ventral striatum, respectively.

We also assessed whether *Lipin1* deficiency had benefit for animal motor function. We examined multiple aspects of cKO/ΔE mice behavior and uncovered two aspects of impairment: a) a broad developmental delay (Figure 5A – 5F) and b) the previously described motor dysfunction (18) (Figure 5G – 5I). We then tested whether either or both was due to Lipin hyperactivity, and thus improved by Lipin deficiency. The answer was negative for all neurodevelopmental measures, including those with (Figure 5C and 5F) and without motor components (like response to auditory stimulus or eye opening; Figure 5D and 5E). In contrast, *abnormal* motor behaviors were sensitive to Lipin deficiency. We detected abnormal forelimb and hindlimb clasping, as well as tremor, in juvenile cKO/ΔE:*Lipin*+/+. The frequency of all three was significantly less in cKO/ΔE:*Lipin*+/− (Figure 5G - 5I) The fact that Lipin deficiency specifically suppressed abnormal movements while other parameters were unaffected (Figure 5A-F) suggests it acts directly on motor circuits, rather than indirectly via improving cKO/ΔE weight or development. Indeed, the significantly reduced frequency of motor dysfunction in cKO/ΔE:*Lipin*+/− compared with cKO/ΔE:*Lipin*+/+ occurred even though the two groups lost unequal numbers of their clinically worst affected individuals (27% vs. 53% lethality, respectively; Figure 2F).

**Figure 5.**
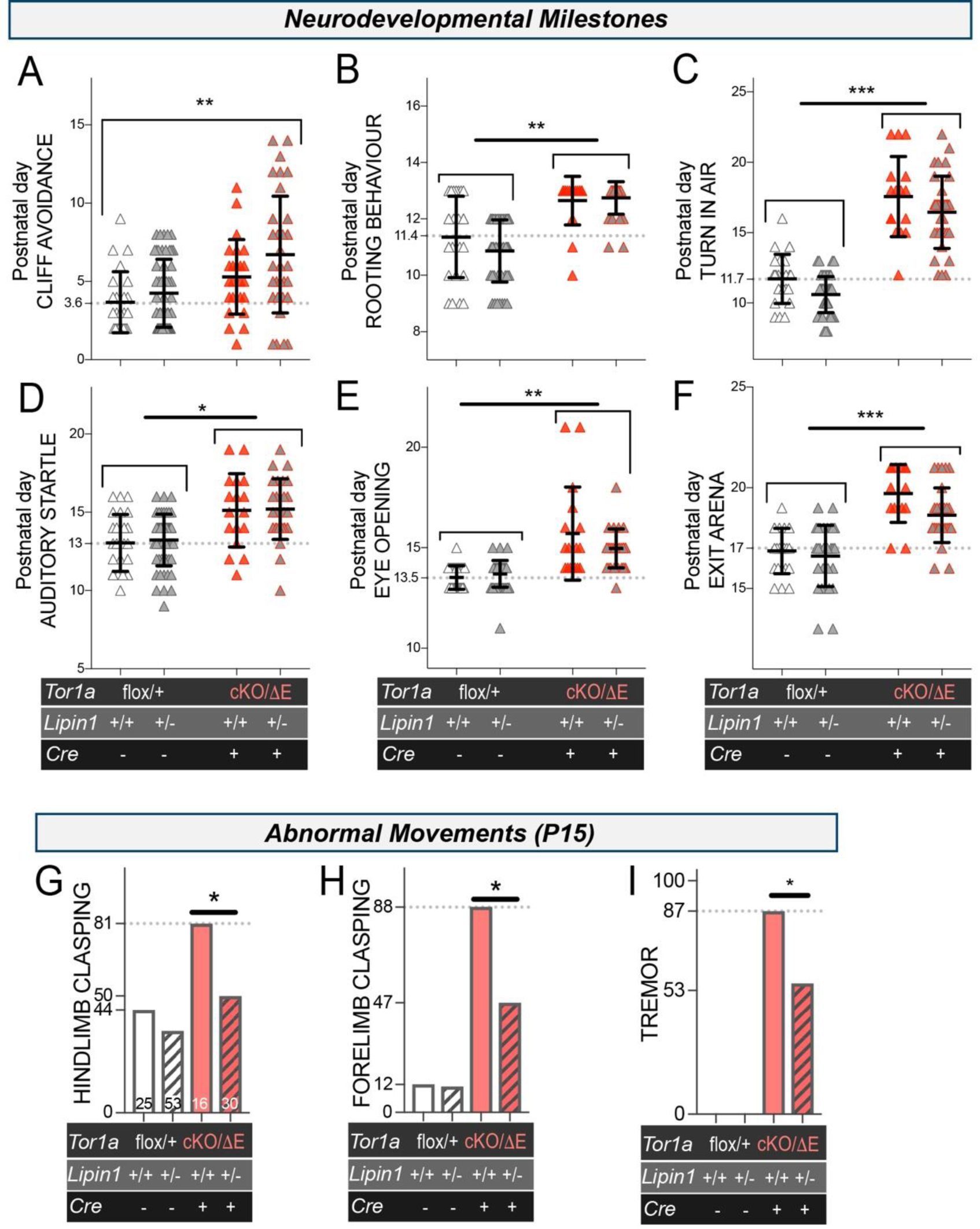
Lipin deficiency rescues movement disorders in cKO/ΔE mice. A – F) Tests that monitor developmental progression (44, 45) detect multiple impairments in cKO/ΔE mice, that are not rescued by *Lipin1*+/−. Animals were examined daily until achieving the milestone/behaviour or being lost from the study. Points show the age that individual mice first performed a behaviour on two consecutive days, with mean and standard deviation also shown for each group. *, **, *** significant difference between both cKO/ΔE groups and both controls (p < 0.05, 0.01, 0.001; One-way ANOVA). G – I) % of P15 animals with motor abnormalities. The % of affected animals is significantly higher for cKO/ΔE:Lipin1+/+ than controls (not indicated), and significantly reduced in cKO/ΔE:Lipin1+/−. * significant difference; p < 0.05. Two-tailed Chi-square test. Values in G show the *n* for each group.

These data establish that Lipin hyperactivity is pathogenic for TorsinA disease symptoms in mice. They also raise questions about why these are neurological given that TorsinA and Lipins are broadly expressed (32, 33). In fact, Lipin activity was normal in TorsinA −/− embryonic liver and heart, as well as in ΔE/ΔE mouse embryonic fibroblasts (MEFs) (Figure 6A); the same animals that had ∼3x Lipin activity in brain (Figure 1D). We investigated why, focusing on homologous *Tor1b*/TorsinB expression that, at least in some circumstances, appears to protect non-neuronal cells against TorsinA loss (7, 34, 35). Indeed, *Tor1b* knock-down (Figure 6B) significantly increased Lipin activity in ΔE/ΔE MEFs, but not in +/+ (Figure 6C). Thus, torsinA and torsinB both regulate Lipin, and torsinB expression can mask the impact of torsinA loss. We then took advantage of this assay to ask whether TorsinA mice accurately model the biochemical landscape of human TorsinA disease. We electroporated human fibroblasts from controls (TorsinA +/+) and patients with TorsinA dystonia (genotype: +/ΔE) with control or TOR1B shRNA (Figure 6D and 6E). This efficiently suppressed TOR1B expression (Figure 6D) concomitant with significantly elevating Lipin activity in patient-derived cells but not in control cells (Figure 6E). Finally, we also examined Lipin activity in lysates of iPSC-derived neurons from two controls and two individuals with TorsinA dystonia that had been differentiated in parallel. The two patient samples had Lipin activity at 1.5 and 2.0 relative to the control (Figure 6F).

**Figure 6.**
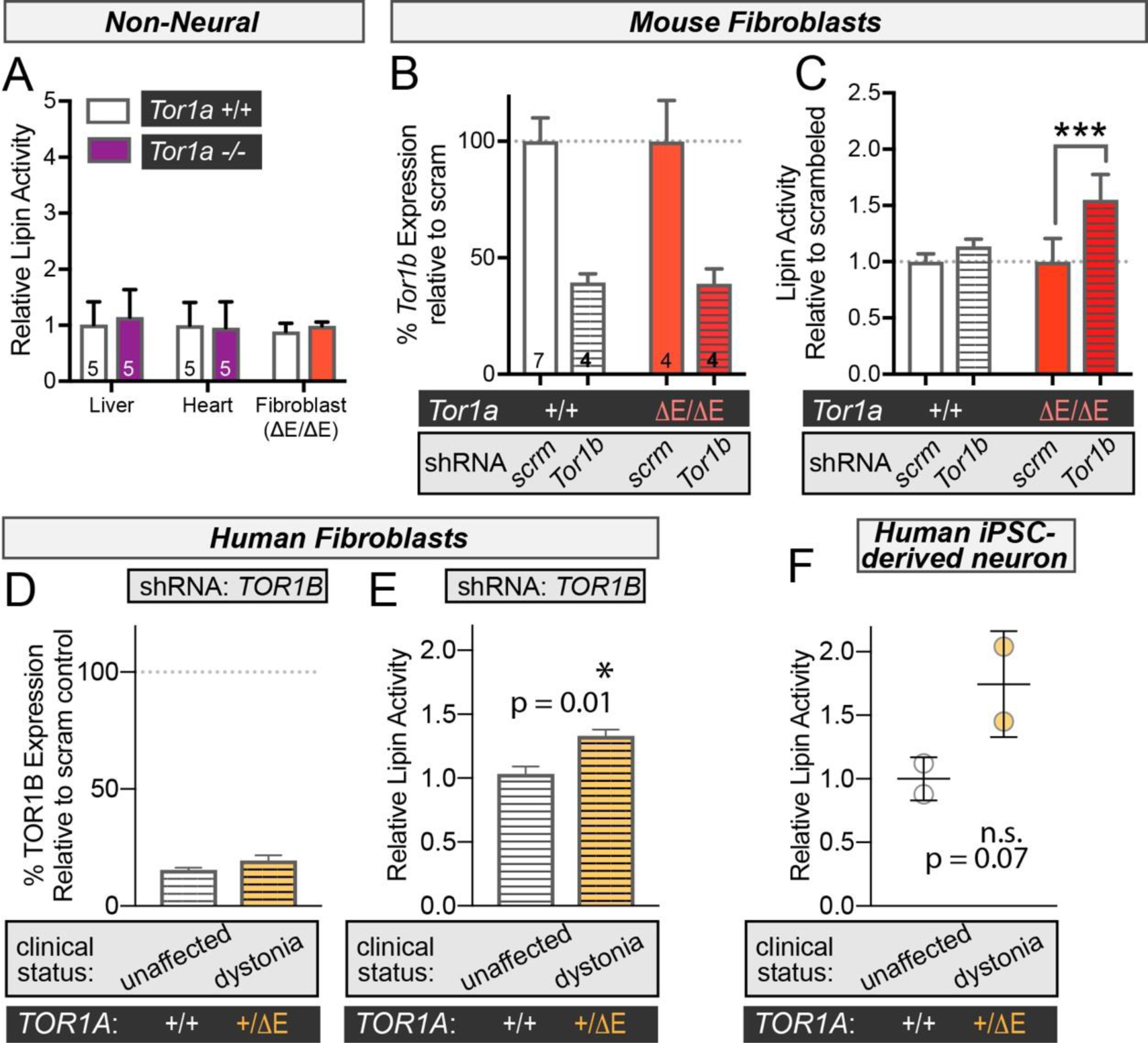
Lipin is abnormal in human TorsinA patient cells. A) Lipin PAP activity in E18.5 tissues and MEFs (n = 3 separate assessments). B – C) *Tor1b* shRNA elevates Lipin activity in mouse ΔE/ΔE fibroblasts. (B) *Tor1b* shRNA similarly reduces *Tor1b* in control and ΔE/ΔE MEFs but (C) specifically elevates Lipin activity in ΔE/ΔE MEFs. Bars show mean +/− SEM. ***, p < 0.001 (2- way ANOVA). *n* indicates separate electroporations of individual cell lines. D – E) Abnormal Lipin regulation in human dystonia patient fibroblast lines. (D) *TOR1B* shRNA efficiently reduces *TOR1B* mRNA in control and patient cells. Bars show mean +/− SD of *TOR1B* expression in control and patient lines relative to that of cells electroporated with the scrambled control. (E) Lipin activity in lysates from human controls and TorsinA dystonia patient fibroblasts treated with *TOR1B* shRNA. Lipin activity is presented relative to that of the same cell line electroporated with the scrambled control. * significant difference between means. p = 0.01; One tailed T-Test (n = 2). F) Lipin activity in iPSC derived neurons from two controls and two individuals with TorsinA dystonia. Lipin activity is presented relative to the control mean, points show values of each individual, and bars show group mean +/− SD. p = 0.07; One tailed T-Test (n = 2).

## Discussion

These data establish a causative relationship between Lipin hyperactivity and the development of TorsinA disease. This conclusion comes from finding excess Lipin activity in neonatal-lethal homozygous TorsinA mice that genetically mimic recessive disease. It comes from finding that Lipin biochemistry is similarly altered in TorsinA patient and mouse fibroblasts, and that Lipin activity appears abnormal in iPSC-derived neurons from TorsinA dystonia patients. Most importantly, it comes from the benefit of Lipin1 deficiency against lethality, neurodegeneration, and abnormal movements of a TorsinA disease model mouse. Indeed, the fact we intervene against behavioral dysfunction underlies why we assign a causative relationship between Lipin and diseases that are defined by their behavioral symptoms.

There are at least two different TorsinA diseases caused by different dosages of TorsinA mutation: +/ΔE causes dominant TorsinA dystonia (1), while ΔE/ΔE (or other mutations) causes broad developmental delay, feeding, speech and motor problems, as well as lethality in some children (2–5). Here we focus on a conditional TorsinA disease mouse model that recapitulates features of both diseases. Firstly, considering its recessive disease symptoms, the mouse has perinatal and juvenile lethality, developmental delay, and tremor. These occur concomitant with neurodegeneration. We show that three of four phenotypes are strongly rescued by partial Lipin deficiency. Thus, Lipin inhibition is a potentially powerful therapeutic approach for children suffering an incurable disease that requires a lifetime of support and care. It is surprising, given the suppression of other phenotypes, that Lipin deficiency has little benefit for slowed development. However, we cannot distinguish whether this truly reflects a Lipin-independent process, or whether it relates to the confound of greater lethality in “non-rescued” cKO/ΔE mice. Alternatively, it could also relate to the fact that Lipin activity remains above normal even in “Lipin-rescued” mice.

The dominant disease of TorsinA dystonia is a movement disorder. Thus, the fact that Lipin deficiency suppresses abnormal movements in cKO/ΔE mice suggests a pathogenic role for Lipin hyperactivity. This is also strongly supported by the fact that Lipin is abnormally regulated in fibroblasts from dystonia patients (TOR1A +/ΔE genotype), and appears elevated in TorsinA dystonia patient iPSC-derived neurons. Moreover, dysfunction of striatal cholinergic neurons is strongly implicated in dominant TorsinA dystonia pathogenesis (31, 36, 37), and we show here that Lipin hyperactivity has a role in their selective vulnerability to TorsinA mutation (19). Thus, these data strongly implicate Lipin hyperactivity in dominant TorsinA dystonia and predict that Lipin inhibition also represents a therapy for these patients.

The data also raise the question of why Lipin hyperactivity is neurotoxic. The mechanism differs from flies where it upregulates triglycerides and suppresses membrane lipid levels (12). The most obvious explanation is that transient and/or locally altered levels of PtdA and/or DAG lipids inappropriately activate signaling cascades, for example impacting mTor and/or protein kinase C that are well characterized responders to PtdA and DAG, respectively (15, 38). In addition, our data firmly establish Lipin regulation as a key role of torsin AAA+ enzymes: it encompasses fly *dTorsin*, human and mouse TorsinA, as well as TorsinB. This places torsins on a list of Lipin regulators that includes insulin and mechano-signaling, amongst others (29, 39, 40). How torsins biochemically execute Lipin regulation also requires additional study, but it is potentially via Lipin phosphorylation given that only phosphorylation-deficient Lipin1 mutants cause cytotoxicity in mammalian cells like occurs when TorsinA is lost.

The most important finding is the therapeutic potential of Lipin inhibition for TorsinA diseases, especially the currently untreatable recessive syndrome. Indeed, an overactive enzyme is amongst the most feasible of druggable targets. The fact we see benefit from relatively mild Lipin inhibition (mediated by the *Lipin1*+/− genotype) is also crucial: this degree of Lipin inhibition is tolerated by mice, as well as humans. Currently there are no small molecules known to inhibit Lipin specifically or with high affinity, and those in experimental use hit other targets with orders of magnitude higher affinity (41, 42). Nevertheless, Lipin has high specificity for a single substrate molecule (PtdA) and belongs to a relatively unstudied and unexploited class of phosphatases (43). This suggests that Lipin inhibition is a realistic druggable target and, given the data presented here, can provide powerful therapeutic benefit for currently incurable conditions that have such severe symptoms to be lethal.

## Acknowledgments

We thank Karolien Billion. and Sergio Hernandez Diaz for technical support, advice from Zsuzsanna Vegh at the KUL Mouse Behavioral Core, Bart de Strooper for manuscript feedback, VIB Bioimaging (particularly Benjamin Pavie) and Nucleomics Facilities, and Lipotype for lipidomics.

## Author contributions

A.C. and R.E.G. were involved in all aspects of project conceptualization, data collection, analysis, supervision, and manuscript preparation. J.F. performed, analyzed and interpreted experiments. S.R, P.S., B.D.G, S.F.G, L.H. and M.M performed experiments. P.V., A.P. and C.K. supervised experiments.

## Data and materials availability

All data is available in the main text or the supplementary materials.

## Methods

### Mouse Husbandry and Tissue Collection

*Lipin1*, *Tor1a* and Nestin-Cre mice are previously described (17, 18, 24, 46) and/ or acquired from Jackson Mice (Bar Harbour, ME). Genotyping was performed as previously described or by TransnetYX (Memphis, TN). The days of embryonic development were defined after assigning the day of vaginal plug detection as E0.5. Embryos were collected from pregnant females after they were euthanized by cervical dislocation. Days of post-natal development were defined with the birthdate as P0. Postnatal animals were permanently identified using the AIMS Pup Tattoo Identification System (Budd Lake, NJ). Tissues were collected from post-natal animals after decapitation (P0 until P14), cervical dislocation (P14 until P35), or CO_2_ inhalation. Tissue destined for biochemical analysis was snap frozen and stored at −80°C until use. Tissues destined for histological analysis were perfused and fixed overnight at 4°C in 4% paraformaldehyde in phosphate buffered saline (PBS). They were then either dehydrated and embedded in paraffin or cryoprotected in 30% sucrose, placed in embedding media, rapidly frozen on dry ice, and stored at −80°C until required.

All mice were housed in the KU Leuven animal facility, fully compliant with European policy on the use of Laboratory Animals. To prevent environmental bias, mice were cohoused independent to genotype. All animal procedures were approved by the Institutional Animal Care and Research Advisory Committee of the KU Leuven (ECD P120/2017) and performed in accordance with the Animal Welfare Committee guidelines of KU Leuven, Belgium.

### Neurological function

P0-P21 pups were weighed each day and tested using adaptions of existing protocols (45, 47). For the cliff aversion test, a mouse was placed on the edge of a smooth-surfaced box with its snout hanging over the edge, and observed for whether it moved away within 30s. Rooting behaviour was examined started at P6, where we examined whether an animal turned its head when brushed with a fine cotton filament. If unresponsive, the test was repeated on the other side of the animal. The ability to right position was scored based on a pup landing on all four paws after being released, oriented legs up, approximately 10cm above a 5cm thick layer of shavings/ cotton wool. The Auditory Startle response was examined with mice placed individually in a fresh cage, and handclapping at a distance of approximately 10cm). A quick involuntary movement of any type was considered positive. Eye opening was scored as complete when both eyes were open. Finally, locomotor and exploratory behaviour milestones were examined after placing a mouse in the center of a 13cm diameter circle drawn in pen on a large piece of card. Animals that moved sufficiency that all four paws exited the circle within 30 seconds.

For tremor, each mouse was observed in its home cage for 20s. Any sign of tremor (mild or severe) was recorded as positive. Forelimb and hindlimb clasping were examined upon suspending a mouse upside down by the tail. A positive score was recorded if the limbs touched. Kyphosis was also recorded if an animal appeared unable to straighten its spine during ambulation or at rest. All tests were performed with all mice that had been cohoused and observers were blind to genotype. While cKO/ΔE animals could be distinguished by reduced weight, this could not discriminate their *Lipin1* genotype. After weaning (P21) we continued to examine animals for overall health, including weighing every fourth day.

### Cell Lines and Electroporation

Mouse embryonic fibroblasts were produced from E13.5 embryos using standard conditions, and transformed by transfection with a plasmid encoding the SV40 large tumor-antigen, followed by > 10 rounds of passaging, and storing in liquid N_2_ until use. shRNA plasmids were electroporated Gene Pulser Xcell™, and cells harvested 72-hours post-transfection for qRT-PCR, protein, and PAP determination. Human fibroblasts were generated by standard protocols. Human fibroblasts were cultured for 6 days prior to electroporation with shRNA plasmids. Electroporation was performed using the Human Adult Dermal Fibroblast Nucleofector™ Kit and the Nucleofector™ II/2b device (Lonza). Both cell lines were harvested 72-hours post-transfection for qRT-PCR, protein, and PAP determination. Hela cells were electroporated with plasmids using Ingenio electroporation kit (SopaChem) and the Nucleofector™ II/2b device (Lonza). Cells were harvested 24h post-electroporation, and the % cell survival quantified using Trypan Blue staining (Thermo Fisher Scientific) and Cell Counter (Biorad).

### Plasmids

ShRNA plasmids used were acquired from Sigma: *Tor1b* (TRCN0000106485), scrambled control (Empty pLKO.1), and *TOR1B* (TRCN0000159398). We used Gibson cloning (NEB) to generate plasmids that express mouse Lipin1 fused at the C-terminus with mCherry. We PCR amplified Lipin1 (lacking a stop codon) and mCherry cDNA sequences and fused these in-frame between the Xho1 and BglII sites of pBudCE4.1 (Invitrogen). Lipin1 sequences were amplified from the following plasmids: pRK5 FLAG 17xS/T->A, catalytic active lipin 1 (Addgene #32007), pRK5 FLAG wildtype, catalytic active lipin 1 (#32005), pRK5 FLAG 17xS/T->A, catalytic dead lipin 1 (#32008), pRK5 FLAG wildtype, catalytic dead lipin 1 (#32005).

### Patient cells, iPSC reprogramming, culturing and differentiation

We included two dystonia patients carrying the +/ΔE mutation in TorsinA (one male and one female) and four healthy control individuals. Skin biopsies for functional assays were taken from each individual. The study was approved by the local ethics committee of the University of Lübeck and all participants gave written informed consent. Two patient skin fibroblast lines were reprogrammed into iPSCs using Sendai virus to deliver the four reprogramming factors OCT4, SOX2, KLF4, and cMYC (CytoTune-iPS 2.0 Sendai Reprogramming Kit, Thermo Fisher Scientific). Two control fibroblast lines were reprogrammed into iPSCs through StemBANCC (48). The iPSC lines were cultured on Matrigel-coated dishes (BD Biosciences) in mTeSR1 medium (STEMCELL Technologies). The direct differentiation of iPSCs into dopaminergic neurons was conducted as described before (49, 50) and neurons were terminally differentiated for 40-50 days.

### PAP Activity

PAP activity was measured according to standard protocols (51, 52) based on the formation of fluorescent DAG from NBD-PA (1-acyl-2-{12-[(7-nitro-2-1,3- benzoxadiazol-4-yl)amino] dodecanoyl}-sn-glycero-3-phosphate ammonium salt; Avanti® Polar lipids, Inc). Snap frozen tissue or cells were homogenized on ice in 50 mM Tris HCl, pH 7.5 containing 0.25 M sucrose, 10 mM 2-mercaptoethanol, 1x PhosSTOP phosphatase inhibitor cocktail (Roche) and 1x EDTA free protease inhibitor cocktail (Sigma). Lysates were centrifuged for 10 min at 1,000 × g at 4 °C, and supernatant transferred. Protein was measured with a modified Bradford assay (Bio-Rad). 60μg aliquots of protein were then incubated for 30 min at 30°C with the same buffer including 2 mM NBD-PA (with 10 mM Triton X-100) and either 0.5 mM MgCl_2_ or 1 mM EDTA. Reactions were terminated by adding 0.4 ml of 0.1 M HCl diluted in methanol, followed by lipid extraction by adding 0.4 ml chloroform and 0.4 ml of 1 M MgCl_2_, centrifuging at 1000 rpm for 10 min at room temperature, and removing the chloroform-soluble lipid fractions to dry under nitrogen. These were then re-solubilized in 30μl chloroform/methanol (1:1) and loaded onto silica gel plates (Merck). PA and DAG were separated by 10-15min of thin layer chromatography (TLC) using chloroform/methanol/H_2_O (62:25:4) as the solvent. Fluorescence intensity was recorded with an ImageQuant LAS 4000 device (Green-RGB, 460nm/534nm). ImageJ was used to measure the density of DAG fluorescence in MgCl_2_ and EDTA conditions. All reactions were performed at least in duplicate, and Lipin PAP activity (arbitrary units) was calculated as the average DAG fluorescence produced by MgCl_2_-containing reactions minus the average DAG fluorescence of EDTA reactions. LPP activity was calculated as the average DAG fluorescence of the EDTA reactions. Values were expressed relative to those of internal control reference samples present on the same TLC plate. Unless otherwise stated, all PAP activity data are presented as relative to the mean wild-type control value.

### RNAseq

#### Sample Preparation and Sequencing

Library preparation, sequencing and statistical data analysis were performed by the VIB Nucleomics Core (www.nucleomics.be). RNA concentration and purity were determined spectrophotometrically using the Nanodrop ND-1000 (Nanodrop Technologies) and RNA integrity was assessed using a Bioanalyser 2100 (Agilent). Per sample, an amount of 500 ng of total RNA was used as input. Using the Illumina TruSeq® Stranded mRNA Sample Prep Kit (Part # 15031048 Rev E. October 2013), poly-A containing mRNA molecules were purified from the total RNA input using poly-T oligo-attached magnetic beads. In a reverse transcription reaction using random primers, RNA was converted into first strand cDNA and subsequently converted into double-stranded cDNA in a second strand cDNA synthesis reaction using DNA Polymerase I and RNAse H. The cDNA fragments were extended with a single ‘A’ base to the 3’ ends of the blunt-ended cDNA fragments after which multiple indexing adapters were ligated introducing different barcodes for each sample. Finally, enrichment PCR (12 cycles) was carried out to enrich those DNA fragments that have adapter molecules on both ends and to amplify the amount of DNA in the library. Sequence-libraries of each sample were equimolarly pooled and sequenced on Illumina NextSeq500 flow-cells with a High Output 75bp kit (Single Read – 1.2pM Pool + 1.89% PhiX v3) at the VIB Nucleomics core (www.nucleomics.be).

#### Data analysis

##### Preprocessing

Low quality ends and adapter sequences were trimmed off from the Illumina reads with FastX 0.0.14 and Cutadapt 1.7.1 (53, 54). Subsequently, small reads (length < 35 bp), polyA-reads (more than 90 % of the bases equal A), ambiguous reads (containing N), low-quality reads (more than 50 % of the bases < Q25) and artifact reads (all but three bases in the read equal one base type) were filtered using using FastX 0.0.14 and ShortRead 1.24.0 (55). With Bowtie2 2.2.4 we identified and removed reads that align to phix_illumina (56).

##### Mapping

The preprocessed reads were aligned with STAR aligner v2.4.1d to the reference genome of *Mus musculus* (GRCm38.73) (57). Default STAR aligner parameter settings were used, except for ‘--outSAMprimaryFlag OneBestScore -- twopassMode Basic --alignIntronMin 50 --alignIntronMax 500000 --outSAMtype BAM SortedByCoordinate’. Using Samtools 1.1, reads with a mapping quality smaller than 20 were removed from the alignments (58).

##### Counting

The number of reads in the alignments that overlap with gene features were counted with featureCounts 1.4.6 (59). Following parameters were chosen: -Q 0 -s 2 -t exon -g gene_id. We removed genes for which all samples had less than 1 count-per-million. Raw counts were further corrected within samples for GC-content and between samples using full quantile normalization, as implemented in the EDASeq package from Bioconductor (60).

##### Differential gene expression

With the EdgeR 3.8.6 package of Bioconductor, a negative binomial generalized linear model (GLM) was fitted against the normalized counts (61). We did not use the normalized counts directly, but worked with offsets. Differential expression was tested for with a GLM likelihood ratio test, also implemented in the EdgeR package. To select the differentially expressed genes, we adopt the criterion that was used during the elaborate MAQC-I study (62) that selected genes based on uncorrected p-value < 0.001 and a fold change >2.

### qRT-PCR

Cells preserved in RNAlater were disrupted and homogenized in RLT buffer using a QIAshredder, followed by RNA isolation according to the manufacturer’s instructions (RNeasy Qiagen Mini Kit; all Qiagen). cDNA was generated using SuperScript IV Reverse Transcriptase (Invitrogen) and 50μM random hexamer priming. cDNA was diluted in nuclease-free water and stored at −20°C until used in qPCR with the SensiFast_TM_ SYBR_®_ No-ROX kit (Bioline) and a Lightcycler® 480/1536 (Roche) under the SYBRGreen standard run protocol. All qPCR runs were performed in duplicate and values for each gene were normalized against three housekeeping genes (HPRT1, PGK1, B2M). Appropriate no-RT and non-template controls were included in each 384-well PCR reaction, and dissociation analysis was performed to confirm reaction specificity. Values were transformed in excel and processed using standard geometric normalization. Primer sequences: *Tor1b* (forward: (CTT GCC ACC GAA GTG ATT; reverse: CAA GGA CAG AGT CAA TGG TTT); *Pgk1* (forward: TTC CCA TGC CTG ACA AGT; reverse: CCC AGC AGA GAT TTG AGT TC); TOR1B (forward: TCA ACG CTT CGG CTC TCA A; reverse: TCA GCG CCT TGA AAA TCA C); PGK1 (forward: CCG CTT TCA TGT GGA GGA AGA AG; reverse: CTC TGT GAG CAG TGC CAA AAG C); HPRT1 (forward: CAT TAT GCT GAG GAT TTG GAA AGG; reverse: CTT GAG CAC ACA GAG GGC TAC A)

### Immunohistochemistry

6µM thick sections of PFA-fixed paraffin-embedded brains were sectioned in a sagittal orientation using an HM355 microtome. Sections were dewaxed and rehydrated using a graded ethanol series. For antigen retrieval, sections were boiled for 20min in sodium citrate buffer (10mM sodium citrate, 0.05% tween-20, pH6.0). Sections were permeabilized and blocked (0.25% Triton X-100 and 10% normal donkey serum in phosphate buffer saline (PBS)) for 1h at room temperature, followed by overnight incubation at 4°C with primary in blocking solution. Primary antibodies used: Anti-choline acetyltransferase (ChAT) (1:50) (AB144P Millipore), Anti-GFAP (1:600) (ab7260), Anti-Cleaved Caspase 3 (Asp 175) (1:400) (Cell signaling).

Sections were washed, incubated for 2h at room temperature with secondary antibody (Rhodamine red anti-goat; Jackson ImmunoResearch) in blocking solution, and washed again. ChAT and Cleaved caspase 3 sections were then counterstained with DAPI (Sigma; 10236276001) and GFAP sections were counterstained with NeuroTrace™ 500/525 Green Fluorescent Nissl Stain (Thermo Fisher Scientific). Sections were mounted using ImmunoHistoMount (ab104131; Abcam). Images were obtained using the Axio Scan.Z1 Slidescanner (Zeiss). Image processing was performed using the ZEN 2.5 software (Zeiss) and quantifications were done on QuPath Version 0.1.2, Java Version 1.8.0_111. Area identification for ChAT and GFAP quantifications was done by a researcher blinded to genotype by comparison with sagittal developmental brain mouse atlas (P14) (http://atlas.brain-map.org), and manually drawing designated areas. Cleaved caspase 3 quantifications were performed across serially sectioned half-hemispheres.

### Statistical analysis

Data are presented as means ± standard deviation (SD) unless stated otherwise. N values are shown in figures as individual points or values, and reflect the number of individual animals unless otherwise stated. No measurements were excluded from any analysis. All biochemical and behavioral analysis of mice were performed by observers blind to genotype. Statistical analysis was performed using Prism 7.04 (GraphPad Software). Post-hoc multiple comparisons following significance in an ANOVA were performed using Dunnett’s test unless otherwise stated. Tests were two-tailed unless otherwise stated.

