## Supplemental Figures for "Inhibition of Lipin lipid phosphatase hyperactivity rescues TorsinA neurological disease"

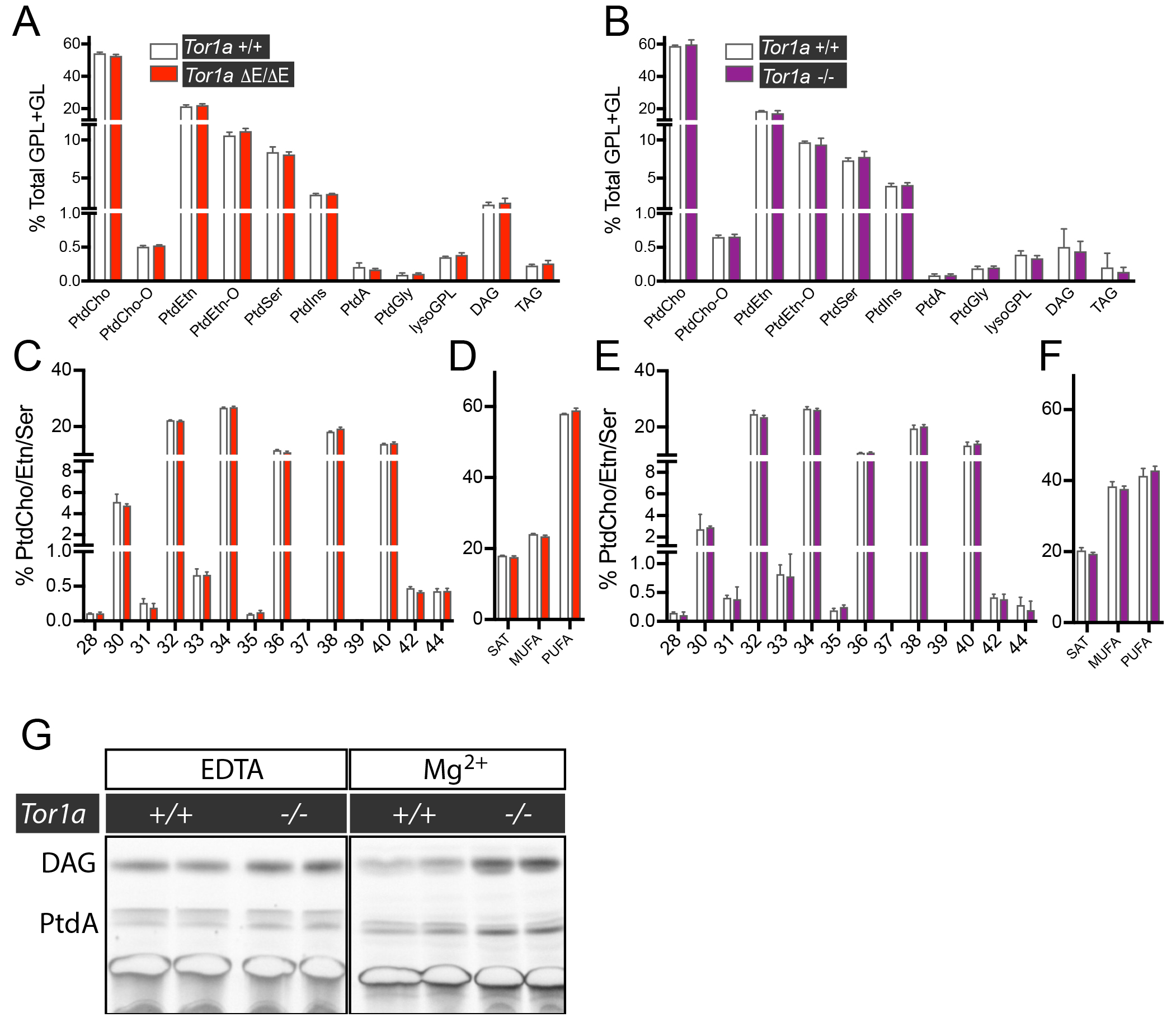

**Supplemental Figure 1. Lipid analyses**

A & B) Relative percentage of glycerophospholipid (GPL) and glycerolipid (GL) classes in the E18.5 spinal cord of littermate (A) TorsinA +/+ (*n* = 2) and ΔE/ΔE (*n* = 3) and (B) +/+ (*n* = 6) and -/- (*n* = 6) mice. No significant differences were detected in the levels of any GPL or GL (multiple T-Tests with two-stage step-up method of Benjamin, Krieger and Yekutieli, with a Q=1%.). –O indicates a lipid ether. (PtdCho) phosphatidylcholine; (PtdEtn) phosphatidylethanolamine; (PtdSer) phosphatidylserine; PtdIns (phosphatidylinositol); (PtdA) phosphatidic acid; (PtdGly) phosphatidyglycerol; lysoGPL (all lyso GPL); DAG (diacylglycerol); (TAG) triacylglycerol.

C & E) Acyl-chain length profile of PtdCho, PtdEtn and PtdSer lipids (~90% of total GPL). Numbers refer to the sum of acyl chain lengths.

D & F) Saturation profile for PtdCho, PtdEtn and PtdSer, including acyl and ether species.

G) Image of fluorescent DAG and PtdA lipids separated by thin layer chromatography. LPP PAP activity is defined as the PtdA to DAG conversion that occurs in the presence of EDTA. Lipin PAP activity is magnesium dependent, and thus calculated from the DAG production (in presence of MgCl_2_), minus the EDTA-resistant fraction.

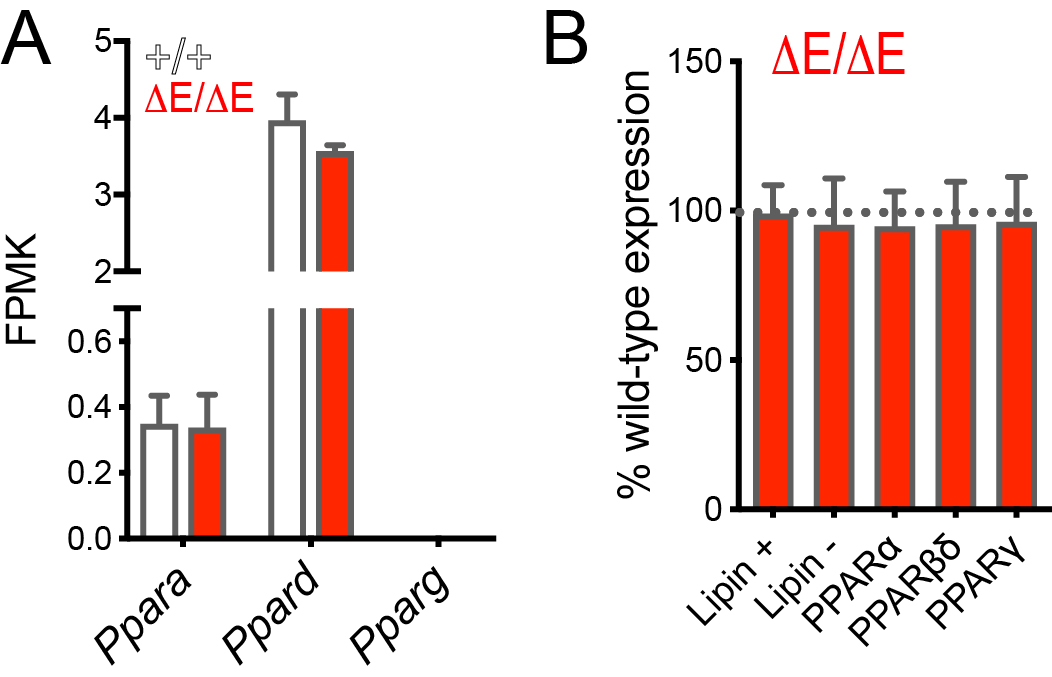

**Supplemental Figure 2. The co-transcriptional function of Lipin is unaffected by TorsinA genotype**

A) PPAR (peroxisome proliferator‐activated receptors) transcription factor expression is unaffected by TorsinA genotype. Bars show mean and SD of FPKM (Fragments Per Kilobase Million) in the E18.5 brain of +/+ (n=12) and ΔE/ΔE (n=6) embryos detected by RNAseq. We did not detect *Pparg* reads. Statistical analysis was by Two-way ANOVA.

B) ΔE/ΔE does not affect expression of genes shown to be induced or suppressed *Lipin1* expression in liver, or targeted by PPAR pathways that are sensitive to the co-transcriptional function of Lipin (1, 2). Gene sets are shown in Supplemental Figure 3. Bars show the percentage change in FPKM between ΔE/ΔE and wild-types from averaging all expression of all genes in each gene set. Gene sets and levels of individual genes are shown in Supplemental Figure 3.

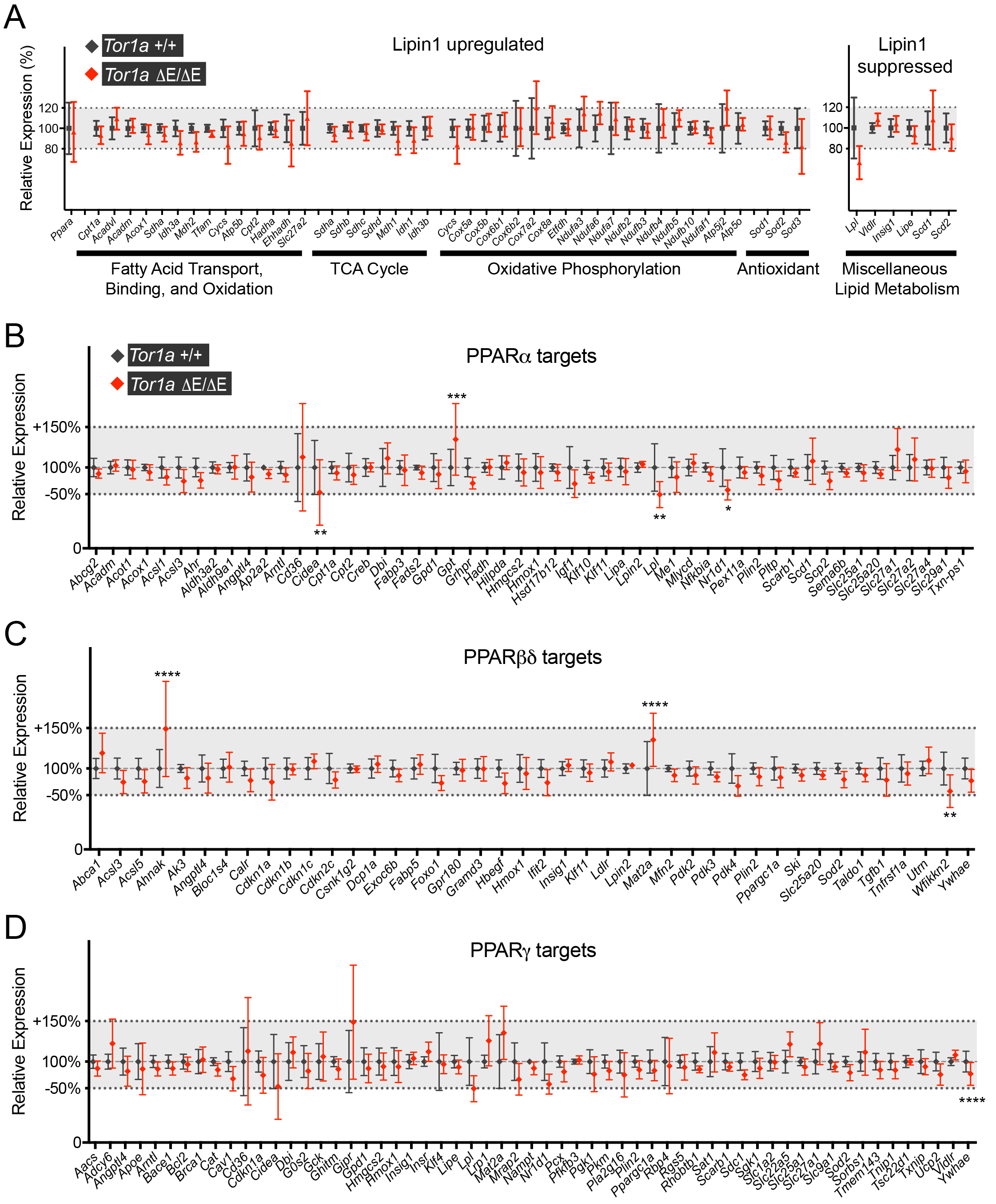

**Supplemental Figure 3. Lipin and PPAR sensitive gene sets**

A) The Lipin+ and Lipin- genes sets that are upregulated and downregulated, respectively, by Lipin1 in liver (1). Symbols show mean +/- SD of FPKM from E18.5 +/+ (n=12) and ΔE/ΔE (n=6) brain normalized to the wild-type mean. No significant effect of genotype was detected (Two-Way ANOVA).

B - D) Expression of PPARα, PPARβδ and PPARγ target genes relative to the wild-type mean. Gene sets were defined using the PPAR gene database (http://www.ppargene.org/index.php). Symbols show the mean +/- SD of FPKM from E18.5 brain of +/+ (n=12) and ΔE/ΔE (n=6), with values of individual embryos first normalized to the wild-type mean. Two-Way ANOVA detected some significantly altered genes within each set. However, none exceeded the 1.5-fold cut-off and 5/8 were present in multiple gene-sets and only reached significance in one analysis.
